## Supplementary Material for "Transcription-driven DNA supercoiling activates bacterial chromatin at a distance"

† Deceased author

✉

**Supplementary Material**

**Supplementary Methods**

**Construction of relevant alleles.**

Strains used in this work were derived from strain MA3409, a derivative of *Salmonella enterica* serovar Typhimurium strain LT2[1] cured for the Gifsy-1 prophage[2]. Generalized transduction was carried out using the high-frequency transducing mutant of phage P22, HT 105/1 *int-201*[3]. Gyrase mutations were introduced by transduction selecting for a linked Tn*10*dCm element[4]. All constructs made by DNA recombineering were obtained in strains harboring λ Red plasmid pKD46[5]. Some of these constructs were described previously[6,7].

Allele Δ[*hilA338*-STM2906]::GFP$^{SF}$-*kan* (also named *hilA*-GFP$^{SF}$ *kan* ΔK28) replaces the right two-thirds of SPI-1 (the 28 Kb segment extending from position 338 of *hilA* to the end of locus STM2906) with a GFP$^{SF}$-*kan* cassette, creating a *hilA*-GFP$^{SF}$ translational fusion (see diagram in Supplementary Fig. 1). The 28 Kb deletion was initially made by inserting an FRT-*kan*-FRT cassette amplified from plasmid pKD13[5] with primers AF93 and AF26; subsequent removal of the cassette by the Flp recombinase left a single FRT site that was used to integrate plasmid pMR6 (which carries a single FRT site fused in frame to the GFP$^{SF}$ orf, plus a *kan* gene for selection) in a second Flp-mediated step, as described[8].

Insertions of a *tetR*-P$^{tet}$ cassette replacing material on the left side of SPI-1 were obtained by two-step scarless recombineering[9]. In the first step, the selectable/counter-selectable *tetR* P$^{tet}$ *ccdB-cat* cassette (amplified with primers AE12 and AH30 from plasmid pNNB5[9]) was used to replace the SPI-1 segment extending from the beginning of *sitA* to the *orgA* gene (allele Δ[*sitA-orgA*]::*tetR*-P$^{tet}$*ccdB-cat*). In the second step, the *ccdB-cat* portion of the construct was replaced by DNA fragments fusing P$^{tet}$ to positions increasingly closer to the *hilD* gene (selecting AHTc-resistance; see Supplementary Figs. 1 and 2). These fragments were generated by "fill-in" PCR of the following oligonucleotide pairs: AG52-AG53 (Δ[*sitA-orgA*]3.0::*tetR*-P$^{tet}$); AG54-AG55 (Δ[*sitA-prgI*]1.6::*tetR*-P$^{tet}$); AH34-AH35 (Δ[*sitA-prgH*]1.2::*tetR*-P$^{tet}$); AH36-AH37 (Δ[*sitA-prgH*]0.8::*tetR*-P$^{tet}$); AH38-AH39 (Δ[*sitA-prgH*]0.6::*tetR*-P$^{tet}$).

Constructs analogous to the ones above, but carrying the full *tetRA* cassette (that is, with P$^{tet}$ fused to *tetA*) were constructed using fragments amplified from the chromosomal DNA of strain MA3397 (which carries a Tn*10*dTc insertion in the Gifsy-1 prophage[2]) selecting tetracycline-resistance. Fragments were made with primer pairs AE12-AF20 (Δ[*sitA-orgA*]3.0::*tetRA*); AE12-AF21 (Δ[*sitA-prgI*]1.6::*tetRA*); AE12-AH23 (Δ[*sitA-prgH*]1.2::*tetRA*); AE12-AH24 (Δ[*sitA-prgH*]0.8::*tetRA*); AE12-AH25 (Δ[*sitA-prgH*]0.6::*tetRA*) (Supplemental Figs 1 and 2).

A strain carrying the *tetRA* cassette in the histidine operon attenuator region (*hisL*) immediately upstream of the Rho-independent transcription terminator (T$^{hisL}$) was constructed using a DNA fragment amplified from MA3397 DNA (see above) with primers AH63 and AH64. This strain (MA14400) was the source of template DNA used to transfer the *tetRA*-T$^{hisL}$ allele to SPI-1. The *tetRA*-T$^{hisL}$ region, amplified with primers AE12-AH67, was transferred by DNA recombineering yielding allele Δ[*sitA-prgH*]1.2::*tetRA*-T$^{hisL}$. A similar step was followed to insert the *tetRA*-T$^{hisL}$ cassette in the *leu* operon promoter region. For this, MA14400 chromosomal DNA was amplified with primers AM62 and AM63. The resulting fragment was introduced into strain MA13387 and recombinants carrying the cassette at the desired position (*leuL*::*tetRA*-T$^{hisL}$) were identified. One of the recombinants (strain MA14606) was used in the present study.

A *tetRA* variant with the *tetA* translation initiation codon changed to AAA was constructed by exchanging the *ccdB-cat* segment in strain MA14332 (above) with a fragment amplified from the MA3397 chromosome (above) with primers AH59 (carrying the desired change at the 5' end of the *tetA*-annealing sequence) and AH61 (selecting for AHTc-resistance).

Insertion of the strong gyrase site (SGS) of phage Mu at a position 0.6 Kb upstream of *hilD* TSS was achieved by first inserting the counter-selectable *tetR*-P$^{tet}$*ccdB-cat* cassette (amplified from plasmid pNNB5 with primers AG44 and AG45) and subsequently exchanging the cassette with a fragment amplified from plasmid pMP310 (which carries the nuB1 variant of Mu SGS[10]) with primers AO27-AO28. The *tetR*-$P^{tet}$*tetA*-$T^{hisL}$ cassette was introduced in the SGS background using a recombineering fragment amplified from the MA14400 chromosome with primers AE12-AO31 (see above). Care was taken in designing primer AO31 to ensure that the 3' boundary of the insertion would lie 189 bp closer to *hilD* than the insertion in the SGS-free context to compensate for the SGS addition (see Supplementary Fig. 5).

When required the *tetA* ORF was exchanged with the *cat* ORF using a recombineering fragment amplified from plasmid pKD3[5] with primers AH55 and AN26.

**Mutagenesis of the SGS.** The segment encompassing the *tetRA*-$T^{hisL}$ cassette and part of the SGS insert of strain MA14726 (*tetR*-$P^{tet}$*tetA*-$T^{hisL}$ SGS *hilA*-$GFP^{SF}$ *kan* ΔK28) was amplified with primers AO37 (Fw primer, annealing near the end of *tetR*) and AO33 (Rv primer, annealing inside the SGS but with the homology interrupted by a randomized 12-nt stretch - beginning 42 nt from the 5' end of the primer - corresponding to the site of gyrase cleavage). The amplified fragment was introduced into strain MA14733 (*tetR*-$P^{tet}$*cat*-$T^{hisL}$ SGS *hilA*-$GFP^{SF}$ *kan* ΔK28 / pKD46) and recombinants were selected on tetracycline-supplemented plates. Colonies showing higher green-fluorescence levels (see Supplementary Fig. 5) were picked and used for PCR and Sanger sequence analysis of the mutagenized region. Performing the same experiment with strain MA14733 as the source of template DNA and strain MA14726 / pKD46 as the recombineering host, allowed isolating SGS mutants in the $P^{tet}$*cat* background.

**5' RACE-Seq Experiments.** 5'RACE-Seq analysis was conducted on three independent RNA preparations from AHTc-treated and untreated cultures of strains MA14692 (Δ[*sitA*-*prgH*]1.2::*tetRA*-$T^{hisL}$) and MA14606 (*leuL*::*tetRA*-$T^{hisL}$). RNA was reverse-transcribed with a mixture of primers AI69 (*hilD*-specific) and AM99 (l*euO*-specifc) in the presence of AI39, the template switching oligonucleotide (TSO). Each of the cDNAs produced (12 samples) was amplified by PCR with primers carrying adapter sequences designed for high throughput sequencing. Oligonucleotide AK47 (which anneals to the TSO sequence) was the common forward primer in all reactions, while a mixture of two oligonucleotides, one annealing in the promoters-proximal region of *hilD*, the other annealing in the promoter-proximal region of *leuO*, and carrying specific index sequences, was used for reverse priming (Supplementary Table 4; amplification program specified in Methods). The PCR products were pooled in equal volumes and the pooled sample was subjected to high throughput sequencing. In the data originating from MA14692, the reads containing the TSO sequence fused to the 5' end of *hilD* were normalized to the reads with the TSO sequence fused to the 5' end of *leuO*. Vice versa for the reads originating for MA14606.

In a separate experiment, the RNA preparations from MA14692 were reverse-transcribed with a mixture of primers AI48 (*prgH*-specific) and AJ33 (*ompA*-specific) in the presence of the TSO. cDNAs was amplified with AJ38 (which anneals to the TSO sequence) as the common forward primer and a single indexed oligonucleotide specific for either *prgH* or *ompA* as the reverse primer. The PCR products were pooled in equal volumes and the pooled sample was subjected to high throughput sequencing. The reads containing the TSO sequence fused to the 5' end of *prgH* were normalized to the reads with the TSO sequence fused to the 5' end of *ompA*.

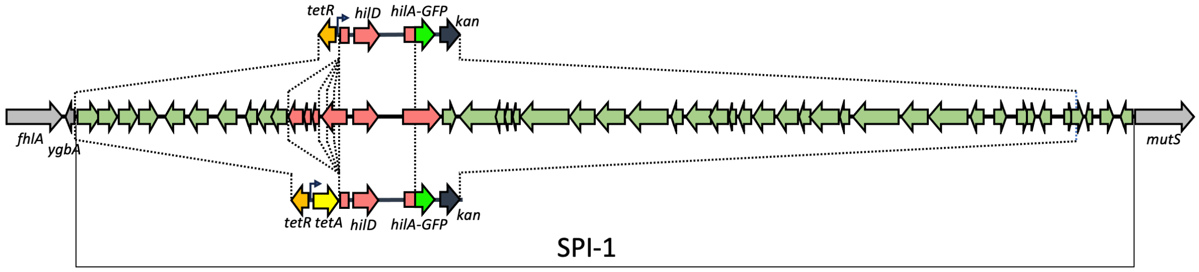


**Supplementary Fig. 1 Schematic diagram showing the gene organization in constructs used in this study**. Insertion of *tetR*-P$^{tet}$ or *tetR*-P$^{tet}$-*tetA* cassette removes a DNA segment extending from the left end of SPI-1 to five different postions between 3 Kb and 0.6 Kb upstream of the *hilD* TSS. Insertions of the *hilA*-GFP$^{SF}$-*kan* cassette removes a 28 Kb portion of SPI-1 extending from inside *hilA* (first 338 bp of the orf) to the end of locus STM2906.

---

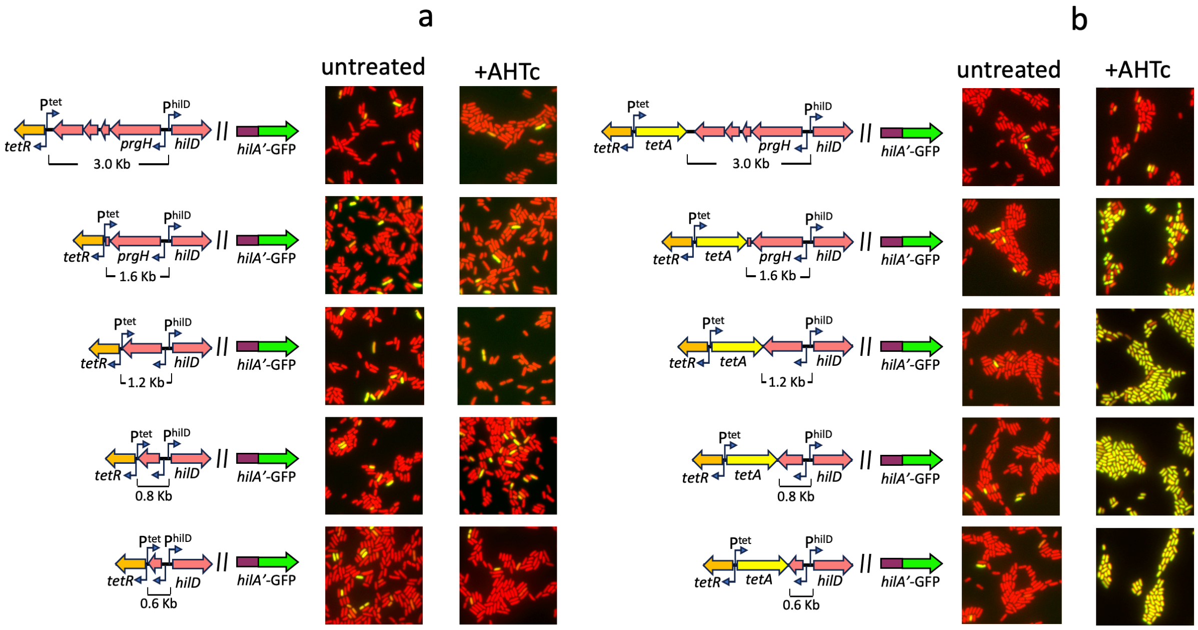


**Supplementary Fig. 2 Comparing the effects of activating the P$^{tet}$ promoter in constructs without (a) or with (b) the *tetA* gene fused to P$^{tet}$**. Corresponding pairs of *tetR*-P$^{tet}$ and *tetR*-P$^{tet}$-*tetA* inserts have identical 3' -flanking sequences. These 3' boundaries fall between 3.0 Kb and 0.6 Kb from the hilD TSS. All strains carry a *hilA*-GFP$^{SF}$ translational fusion and a constitutively expressed mCherry fusion to the P$^{tac}$ promoter. Strains were grown at 37°C to early stationary phase and cells visualized by fluorescence microscopy under 100 x magnification. Representative areas of the microscope field are shown.

---

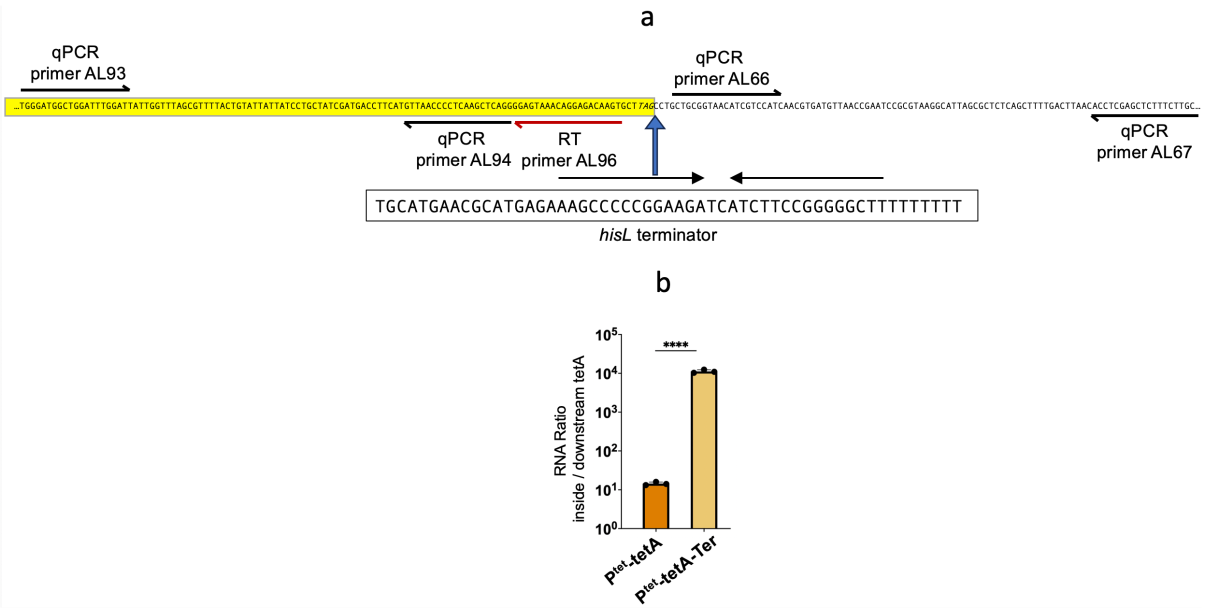


**Supplementary Fig. 3 Assessing the efficacy of the *hisL* terminator. a**. Sequence of the *tetA*-*prgH* boundary (5'-3') showing the site of insertion site of the *hisL* terminator (immediately 3' to the *tetA* stop codon; *tetA* sequence shaded in yellow). Also shown are the positions of RT-qPCR primers used for the quantification of the RNA. **b**. Quantification of the RNA gradient. RNA from cultures grown in the presence of AHTc (0.4 µg/ml) was reverse-transcribed in two separate reactions, one with primer AL96 and primer AJ33 (*ompA*-specific, used for normalization), the other with primers AI11 and AJ33 (RT primer AI11, not shown in the diagram, anneals 38 bp downstream of qPCR reverse primer AL67). The cDNAs produced were amplified by qPCR with primer pairs AL93-AL94 and AL66-AL67, respectively. The data shown represent the ratios between the normalized Ct values obtained in the two amplifications in strains MA14694 ($P^{tet}$-*tetA*) and MA14692 ($P^{tet}$-*tetA*-Ter). The data originate from 3 independent experiments. Significance was determined by the unpaired two-tailed Student's t test (****, P ≤ 0.0001).

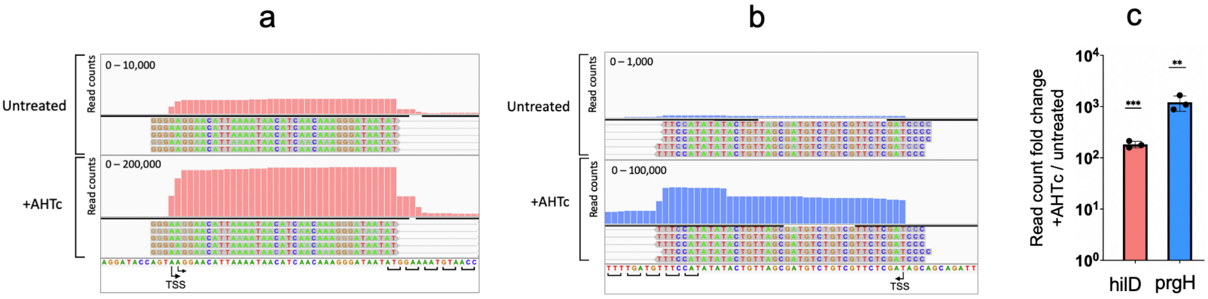


**Supplementary Fig. 4 5'RACE-Seq analysis of *hilD* and *prgH* mRNAs.** Total RNA was extracted from cultures of strain MA14692 (Δ[*sitA*-*prgH*1.2]::*tetRA*-$T^{hisL}$) grown at 37°C in the presence or absence of AHTc (0.4 µM). RNA was reverse-transcribed with either a mixture of primers AI69 (*hilD*) and AM99 (*leuO*) (**a**) or AI48 (*prgH*) and AJ33 (*ompA*) (**b**) in the presence of template-switching oligonucleotide (TSO) AI39. The resulting cDNA was subjected to semi-quantitative PCR amplification and high-throughput sequencing as described in Supplementary Methods. Sequence reads were trimmed to remove the TSO sequence up to the terminal 3 Gs. **a** and **b**. Representative Integrative Genomes Viewer (IGV) snapshots of reads in the *hilD* and *prgH* promoter regions. **c**. Semi-quantitative assessment of the abondance of *hilD* and *prgH* mRNA 5' ends. The counts of reads with the TSO sequence fused to *hilD* or *prgH* 5' ends were normalized to the counts of reads in which the TSO was fused to *leuO* and *ompA* 5' ends, respectively. Results shown are from three independent experiments. Significance was determined by unpaired two-tailed Student's t tests (**, P ≤ 0.01; ***, P ≤ 0.001).

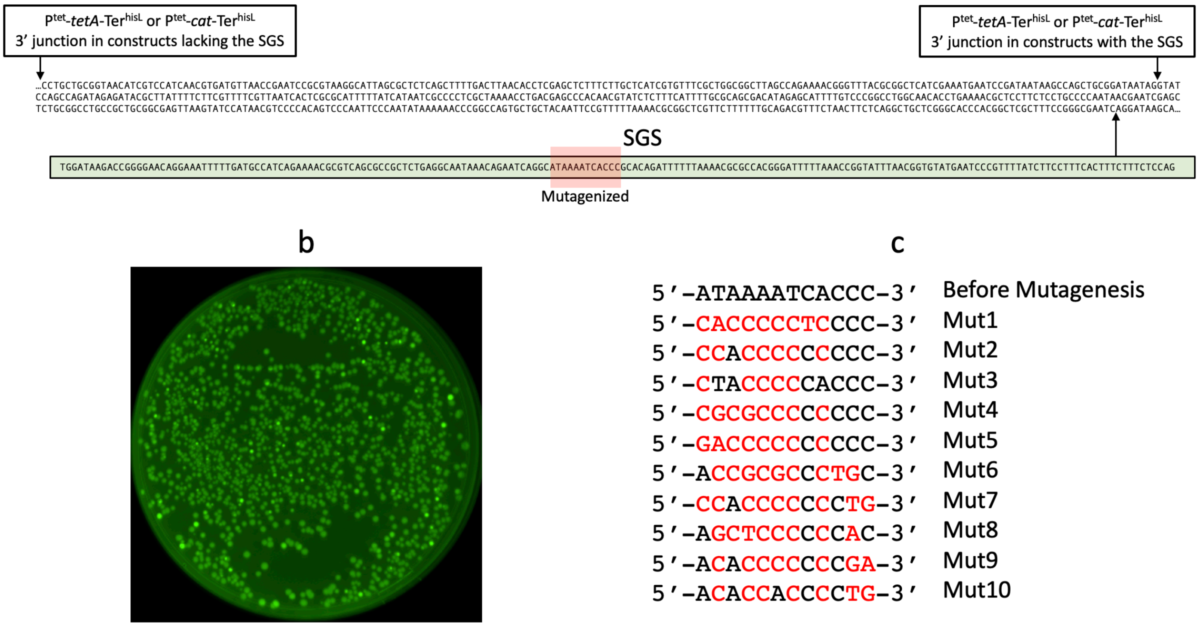


**Supplementary Fig. 5** Experiments with the strong gyrase site (SGS). **a**. Structure of the constructs carrying the SGS. The upward-pointing arrow marks the site of insertion of SGS-containing fragment (189 bp, green-shaded box, oriented as it appears in the genome of bacteriophage Mu). Two downward-pointing arrows indicate the position of the 3' boundaries of the P$^{tet}$*tetA*-Ter insertions. In constructs carrying the SGS insert, this boundary is moved 189 bp downstream to compensate for the SGS addition and maintain the overall distance between the *tetA* cassette and *hilD* unchanged. A 12-bp sequence (encompassing the gyrase cleavage site) subjected to random mutagenesis is shaded in light red. **b**. Representative image of a plate containing colonies from the mutagenic experiment (taken with a Biorad ChemiDoc Touch imaging system under green fluorescence detection conditions). **c**. Sequences of the mutagenized region from 10 clones showing higher fluorescence. Red lettering denotes difference from the original Mu nuB1 sequence[10].

---

### Supplementary Table 1. Relevant strains used in this work

| Strain | Genotype | Reference |
|---|---|---|
| LT2 | wild-type | [1] |
| MA1249 | hisC9968::MudJ gyrB1820 zid-6782::Tn10dCm | [4] |
| MA1564 | din-1001::MudJ gyrA208 zej-6790::Tn10dCm | [4] |
| MA3397 | zfh-8157::Tn10dTc sbcE21 | [2] |
| MA3409 | Δ[Gifsy-1] | [2] |
| MA12803 | Δ[Gifsy-1] hns-3xFLAG FRT-kan-FRT | [7] |
| MA13387 | Δ[Gifsy-1] leuO-lacZY kan | [6] |
| MA14067 | Δ[Gifsy-1] Δ[Gifsy-2] P$^{BAD}$gtgR-P$^{G2R}$nusG-cat Δ[nusG]::aadA hilA338-STM2906]::FRT-kan-FRT | [7] |
| MA14082 | Δ[Gifsy-1] Δ[Gifsy-2] P$^{BAD}$gtgR-P$^{G2R}$nusG-cat Δ[nusG]::aadA Δ[hilA338-STM2906]::GFP$^{SF}$-kan | [7] |
| MA14315 | Δ[Gifsy-1] Δ[his]::P$^{Tac}$mCherry Δ[hilA338-STM2906]::GFP$^{SF}$-kan | [7] |
| MA14320 | Δ[Gifsy-1] Δ[his]::P$^{Tac}$mCherry Δ[sitA-orgA]3.0::tetR-P$^{tet}$tetA Δ[hilA338-STM2906]::GFP$^{SF}$-kan | this work |
| MA14321 | Δ[Gifsy-1] Δ[his]::P$^{Tac}$mCherry Δ[sitA-prgI]1.6::tetR-P$^{tet}$tetA Δ[hilA338-STM2906]::GFP$^{SF}$-kan | this work |
| MA14326 | Δ[Gifsy-1] Δ[Gifsy-2] P$^{BAD}$gtgR-P$^{G2R}$nusG-cat Δ[nusG]::aadA Δ[his]::P$^{Tac}$mCherry Δ[sitA-prgH]1.2::tetR-P$^{tet}$tetA Δ[hilA338-STM2906]::GFP$^{SF}$-kan | this work |
| MA14327 | Δ[Gifsy-1] Δ[Gifsy-2] P$^{BAD}$gtgR-P$^{G2R}$nusG-cat Δ[nusG]::aadA Δ[his]::P$^{Tac}$mCherry Δ[sitA-prgH]0.8::tetR-P$^{tet}$tetA Δ[hilA338-STM2906]::GFP$^{SF}$-kan | this work |
| MA14328 | Δ[Gifsy-1] Δ[Gifsy-2] P$^{BAD}$gtgR-P$^{G2R}$nusG-cat Δ[nusG]::aadA Δ[his]::P$^{Tac}$mCherry Δ[sitA-prgH]0.6::tetR-P$^{tet}$tetA Δ[hilA338-STM2906]::GFP$^{SF}$-kan | this work |
| MA14332 | Δ[Gifsy-1] Δ[his]::P$^{Tac}$mCherry Δ[sitA-orgA]3.0::tetR-P$^{tet}$ccdB cat Δ[hilA338-STM2906]::GFP$^{SF}$-kan | this work |
| MA14339 | Δ[Gifsy-1] Δ[his]::P$^{Tac}$mCherry Δ[sitA-prgH]0.6::tetR-P$^{tet}$ Δ[hilA338-STM2906]::GFP$^{SF}$-kan | this work |
| MA14340 | Δ[Gifsy-1] Δ[his]::P$^{Tac}$mCherry Δ[sitA-prgH]0.8::tetR-P$^{tet}$ Δ[hilA338-STM2906]::GFP$^{SF}$-kan | this work |
| MA14341 | Δ[Gifsy-1] Δ[his]::P$^{Tac}$mCherry Δ[sitA-prgH]1.2::tetR-P$^{tet}$ Δ[hilA338-STM2906]::GFP$^{SF}$-kan | this work |
| MA14348 | Δ[Gifsy-1] Δ[his]::P$^{Tac}$mCherry Δ[sitA-orgA]3.0::tetR-P$^{tet}$ Δ[hilA338-STM2906]::GFP$^{SF}$-kan | this work |
| MA14349 | Δ[Gifsy-1] Δ[his]::P$^{Tac}$mCherry Δ[sitA-prgI]1.6::tetR-P$^{tet}$ Δ[hilA338-STM2906]::GFP$^{SF}$-kan | this work |
| MA14400 | Δ[Gifsy-1] hisL::tetR-P$^{tet}$tetA | this work |
| MA14403 | Δ[Gifsy-1] Δ[his]::P$^{Tac}$mCherry Δ[sitA-prgH]1.2::tetR-P$^{tet}$tetA$^{AAA}$ Δ[hilA338-STM2906]::GFP$^{SF}$-kan | this work |
| MA14443 | Δ[Gifsy-1] hns-3xFLAG FRT Δ[sitA-prgH]1.2::tetR-P$^{tet}$tetA-T$^{hisL}$ | this work |
| MA14533 | Δ[Gifsy-1] Δ[his]::P$^{Tac}$mCherry prgH(0.6)::tetR-P$^{tet}$ccdB cat Δ[hilA338-STM2906]::GFP$^{SF}$-kan | this work |
| MA14606 | Δ[Gifsy-1] leuL::tetR-P$^{tet}$tetA-T$^{hisL}$ leuO-lacZY kan | this work |
| MA14692 | Δ[Gifsy-1] Δ[his]::P$^{Tac}$mCherry Δ[sitA-prgH]1.2::tetR-P$^{tet}$tetA-T$^{hisL}$ Δ[hilA338-STM2906]::GFP$^{SF}$-kan | this work |
| MA14694 | Δ[Gifsy-1] Δ[his]::P$^{Tac}$mCherry Δ[sitA-prgH]1.2::tetR-P$^{tet}$tetA Δ[hilA338-STM2906]::GFP$^{SF}$-kan | this work |
| MA14696 | Δ[Gifsy-1] Δ[his]::P$^{Tac}$mCherry Δ[sitA-prgH]1.2::tetR-P$^{tet}$cat-T$^{hisL}$ Δ[hilA338-STM2906]::GFP$^{SF}$-kan | this work |
| MA14724 | Δ[Gifsy-1] Δ[his]::P$^{Tac}$mCherry prgH(0.6)::SGS(189) Δ[hilA338-STM2906]::GFP$^{SF}$-kan | this work |
| MA14726 | Δ[Gifsy-1] Δ[his]::P$^{Tac}$mCherry Δ[sitA-prgH]1.0::tetR-P$^{tet}$tetA-T$^{hisL}$ prgH(0.6)::SGS(189) Δ[hilA338-STM2906]::GFP$^{SF}$-kan | this work |
| MA14732 | Δ[Gifsy-1] Δ[his]::P$^{Tac}$mCherry Δ[sitA-prgH]1.0::tetR-P$^{tet}$cat-T$^{hisL}$ prgH(0.6)::SGS(189) Δ[hilA338-STM2906]::GFP$^{SF}$-kan | this work |
| MA14755 | Δ[Gifsy-1] Δ[his]::P$^{Tac}$mCherry Δ[sitA-prgH]1.0::tetR-P$^{tet}$cat-T$^{hisL}$ prgH(0.6)::SGS-mut1(189) Δ[hilA338-STM2906]::GFP$^{SF}$-kan | this work |
| MA14793 | Δ[Gifsy-1] Δ[his]::P$^{Tac}$mCherry Δ[sitA-prgH]1.0::tetR-P$^{tet}$tetA-T$^{hisL}$ prgH(0.6)::SGS-mut2(189) Δ[hilA338-STM2906]::GFP$^{SF}$-kan | this work |
| MA14804 | Δ[Gifsy-1] Δ[his]::P$^{Tac}$mCherry Δ[sitA-prgH]1.0::tetR-P$^{tet}$tetA-T$^{hisL}$ prgH(0.6)::SGS(189) gyrA208 zej-6790::Tn10dCm Δ[hilA338-STM2906]::GFP$^{SF}$-kan | this work |
| MA14805 | Δ[Gifsy-1] Δ[his]::P$^{Tac}$mCherry Δ[sitA-prgH]1.0::tetR-P$^{tet}$tetA-T$^{hisL}$ prgH(0.6)::SGS(189) gyrA$^{+}$ zej-6790::Tn10dCm Δ[hilA338-STM2906]::GFP$^{SF}$-kan | this work |
| MA14812 | Δ[Gifsy-1] Δ[his]::P$^{Tac}$mCherry Δ[sitA-prgH]1.0::tetR-P$^{tet}$tetA-T$^{hisL}$ prgH(0.6)::SGS(189) gyrB1820 zid-6782::Tn10dCm Δ[hilA338-STM2906]::GFP$^{SF}$-kan | this work |
| MA14813 | Δ[Gifsy-1] Δ[his]::P$^{Tac}$mCherry Δ[sitA-prgH]1.0::tetR-P$^{tet}$tetA-T$^{hisL}$ prgH(0.6)::SGS(189) gyrB$^{+}$ zid-6782::Tn10dCm Δ[hilA338-STM2906]::GFP$^{SF}$-kan | this work |

### Supplementary Table 2. DNA oligonucleotides used for λ red recombineering

| Name | Sequence (5'-3')[a] |
|---|---|
| AE12 | GCAAAGGCTATATTCGATGATTAATTAACCACATTGTTGCGAGTTAAGACCCACTTTCACATTTA |
| AF20 | GGCGGCAAATGAGTTAATACTGGCGGCATGGCGGCTTAAGAACGGCTAAGCACTTGTCTCCTGTTTA |
| AF21 | GATACGGGCGTTGATAATCTACAAACGCAGGTAACAGAGGCGCTGGCTAAGCACTTGTCTCCTGTTTA |
| AF26 | CCGATGGCAAAGCAAAAGTTTAAAATCAACATCGGAGCGGCAGCTGTAGGCTGGAGCTGCTTC |
| AF93 | GTACGGACAGGGCTATCGGTTTAATCGTCCGGTCGTAGTGGTGTCGATCCGTCGACCTGCAGTTC |
| AG44 | CTTCTCAGGCTGCTCGGGCACCCACGGCTCGCTTTCCGGGCGAATCTTAAGACCCACTTTCACATTTA |
| AG45 | CGGTGCAATTAAATACGCCAATACAGGTCGGTGAATTGCTTATCCTGGTCATTGGCATGTTCAA |
| AG52 | CATTGATAGAGTTATTTTACCACTCCCTATCAGTGATAGAGAAAAGTGCCGTTCTTAAGCCG |
| AG53 | ACCCGCCGCCAGGGCGGCGGCAAATGAGTTAATACTGGCGGCATGGCGGCTTAAGAACGGCAC |
| AG54 | CATTGATAGAGTTATTTTACCACTCCCTATCAGTGATAGAGAAAAGTGCCAGCGCCTCTGTTACC |
| AG55 | ACGTCTCAGCAAAATTTGATACGGGCGTTGATAATCTACAAACGCAGGTAACAGAGGCGCTGGC |
| AH23 | GGATTCGGTTAACATCACGTTGATGGACGATGTTACCGCAGCAGGCTAAGCACTTGTCTCCTGTTTA |
| AH24 | GGGCAGGAGAAGGAGCGTTTTCAGGTGTTGCCAGGCCGGGACACTAAGCACTTGTCTCCTGTTTA |
| AH25 | CGGTGCAATTAAATACGCCAATACAGGTCGGTGAATTGCTTATCCTCTAAGCACTTGTCTCCTGTTTA |
| AH30 | GGCGGCAAATGAGTTAATACTGGCGGCATGGCGGCTTAAGAACGGTCATTGGCATGTTCAA |
| AH34 | CATTGATAGAGTTATTTTACCACTCCCTATCAGTGATAGAGAAAAGTGCCTGCTGCG |
| AH35 | GGATTCGGTTAACATCACGTTGATGGACGATGTTACCGCAGCAGGCACTTTTCTC |
| AH36 | CATTGATAGAGTTATTTTACCACTCCCTATCAGTGATAGAGAAAAGTGTCCCGGCC |
| AH37 | GGGGCAGGAGAAGGAGCGTTTTCAGGTGTTGCCAGGCCGGGACACTTTTCTC |
| AH38 | CATTGATAGAGTTATTTTACCACTCCCTATCAGTGATAGAGAAAAGTGGATAAGCAATTCAC |
| AH39 | CGGTGCAATTAAATACGCCAATACAGGTCGGTGAATTGCTTATCCACTTTTCTC |
| AH55 | GAGTTATTTTACCACTCCCTATCAGTGATAGAGAAAAGTGAAATGGAGAAAAAAATCACTGGATATACC |
| AH59 | GATAGAGTTATTTTACCACTCCCTATCAGTGATAGAGAAAAGTGAAAAAAATAGTTCGACAAAGATCG |
| AH61 | CGGTTAACATCACGTTGATGGACGATGTTACCGCAGCAGGCTAAGCACTTGTCTCCTGTTTACTC |
| AH63 | GTGACTGACAATTCATACTATAAATGCGAAATGAAAAAAGCGCTTAAGACCCACTTTCACATTTA |
| AH64 | GGAAGATGATCTTCCGGGGGCTTTCTCATGCGTTCATGCACTAAGCACTTGTCTCCTGTTTA |
| AH67 | GGATTCGGTTAACATCACGTTGATGGACGATGTTACCGCAGCAGGTAACATTCTGCGTTCCTCTTTA |
| AM62 | GCGTTGAGTAGTAGTAGCCCAGTGAAACGAACGATATGTGACATTTAAGACCCACTTTCACATTTAA |
| AM63 | GTACGTATCCGGATATCGTCAACAAAATGCAATGGCGACAGTAACATTCTGCGTTCCTCTTTA |
| AN26 | CCCGGAAGATGATCTTCCGGGGGCTTTCTCATGCGTTCATGCATTACGCCCCGCCCT |
| AO27 | CTAACTTCTCAGGCTGCTCGGGCACCCACGGCTCGCTTTCCGGGCGAATCTGGATAAGACCGGGGAACA |
| AO28 | CGGTGCAATTAAATACGCCAATACAGGTCGGTGAATTGCTTATCCTCTGGAGAAAGAAAGTGAAAGGAAG |
| AO31 | GATTAACGAAAACGAAGAAAATAAGCGTATCTCTATCTGGCTGGATACTAACATTCTGCGTTCCTCTTTA |
| AO33 | CCGGTTTAAAAATCCCGTGGCGCGTTTTAAAAAATCTGTGCNNNNNNNNNNNNGCCTGATTCTGTTTATTGCCTC |
| AO37 | CCGACCTCATTAAGCAGCTCTAATGCGCTGTTAATCACTTTAC |

[a] Red lettering denotes sequences annealing to template DNA (plasmid, chromosomal, or primer DNA in fill-in PCR).

**Supplementary Table 3. DNA oligonucleotides used as primers for reverse transcription and qPCR**

| Name | Sequence (5’-3’) |
|---|---|
| AI11 | CATTTCGATGAGCCGCGTAA |
| AI48 | ACCGACCTGTATTGGCGTAT |
| AI50 | CGCATCCGTATCCACCTGG |
| AI51 | GACAGGCCGAACACTCTTTG |
| AI62 | TGCCGCAGATAACTTACAGAA |
| AI63 | GTCAGTTTACCGCTCCGAAA |
| AI69 | CAACATCCCAGGTTCGTCAC |
| AJ32 | CGTTGGAGATATTCATGGCGT |
| AJ33 | ACCAGTCGTAGCCCATTTCA |
| AJ37 | GAGACCAGCCCAGTTTAGCA |
| AL66 | CTGCGGTAACATCGTCCATC |
| AL67 | GCAAGAAAGAGCTCGAGGTG |
| AL93 | TGGGATGGCTGGATTTGGAT |
| AL94 | CCTGAGCTTGAGGGGTTAAC |
| AL96 | GCACTTGTCTCCTGTTTACTCC |
| AM99 | CGGCTGAATTCCTCGTCCAT |
| AN20 | GCAAACCACAGCTTCGCAT |
| AN21 | CGACATTCCCAGCGTGTG |

**Supplementary Table 4. DNA oligonucleotides used as primers for 5’ RACE-Seq**

| Name | Sequence (5’-3’)[a] | Anneals to |
|---|---|---|
| AI39 | GCTAATCATTGCAAGCAGTGGTATCAACGCAGAGTACATrGrGrG (TSO) | |
| AJ38 | AATGATACGGCGACCACCGAGATCTACACTCTTTCCCTACACGACGCTCTTCCGATCTCATTGCAAGCAGTGGTATCAAC | TSO |
| AK47 | AATGATACGGCGACCACCGAGATCTACACTCTTTCCCTACACGACGCTCTTCCGATCTGCAGTGGTATCAACGCAGAG | TSO |
| AL01 | CAAGCAGAAGACGGCATACGAGAAAGGAATGTGACTGGAGTTCAGACGTGTGCTCTTCCGATCTGAGACCAGCCCAGTTTAGCA | ompA |
| AL02 | CAAGCAGAAGACGGCATACGAGAATAGTTGGTGACTGGAGTTCAGACGTGTGCTCTTCCGATCTGAGACCAGCCCAGTTTAGCA | ompA |
| AL13 | CAAGCAGAAGACGGCATACGAGAAAGGAATGTGACTGGAGTTCAGACGTGTGCTCTTCCGATCTCGCATCCGTATCCACCTGG | prgH |
| AL14 | CAAGCAGAAGACGGCATACGAGAATAGTTGGTGACTGGAGTTCAGACGTGTGCTCTTCCGATCTCGCATCCGTATCCACCTGG | prgH |
| AL17 | CAAGCAGAAGACGGCATACGAGAAGTCGTCGTGACTGGAGTTCAGACGTGTGCTCTTCCGATCTGAGACCAGCCCAGTTTAGCA | ompA |
| AL18 | CAAGCAGAAGACGGCATACGAGAACTTCGAGTGACTGGAGTTCAGACGTGTGCTCTTCCGATCTGAGACCAGCCCAGTTTAGCA | ompA |
| AL39 | CAAGCAGAAGACGGCATACGAGAAGTCGTCGTGACTGGAGTTCAGACGTGTGCTCTTCCGATCTCGCATCCGTATCCACCTGG | prgH |
| AL40 | CAAGCAGAAGACGGCATACGAGAACTTCGAGTGACTGGAGTTCAGACGTGTGCTCTTCCGATCTCGCATCCGTATCCACCTGG | prgH |
| AN72 | CAAGCAGAAGACGGCATACGAGAATGGTCAGTGACTGGAGTTCAGACGTGTGCTCTTCCGATCTGCTGCCGGGTATTTGTCAAA | hilD |
| AN73 | CAAGCAGAAGACGGCATACGAGAACACTGTGTGACTGGAGTTCAGACGTGTGCTCTTCCGATCTGCTGCCGGGTATTTGTCAAA | hilD |
| AN74 | CAAGCAGAAGACGGCATACGAGAACTGATCGTGACTGGAGTTCAGACGTGTGCTCTTCCGATCTGCTGCCGGGTATTTGTCAAA | hilD |
| AN75 | CAAGCAGAAGACGGCATACGAGAATTTCACGTGACTGGAGTTCAGACGTGTGCTCTTCCGATCTGCTGCCGGGTATTTGTCAAA | hilD |
| AN76 | CAAGCAGAAGACGGCATACGAGAACCACTCGTGACTGGAGTTCAGACGTGTGCTCTTCCGATCTGCTGCCGGGTATTTGTCAAA | hilD |
| AN77 | CAAGCAGAAGACGGCATACGAGAAATCAGTGTGACTGGAGTTCAGACGTGTGCTCTTCCGATCTGCTGCCGGGTATTTGTCAAA | hilD |
| AN78 | CAAGCAGAAGACGGCATACGAGAAAGGAATGTGACTGGAGTTCAGACGTGTGCTCTTCCGATCTGCTGCCGGGTATTTGTCAAA | hilD |
| AN79 | CAAGCAGAAGACGGCATACGAGAATAGTTGGTGACTGGAGTTCAGACGTGTGCTCTTCCGATCTGCTGCCGGGTATTTGTCAAA | hilD |
| AN80 | CAAGCAGAAGACGGCATACGAGAAAAAATGGTGACTGGAGTTCAGACGTGTGCTCTTCCGATCTGCTGCCGGGTATTTGTCAAA | hilD |
| AN81 | CAAGCAGAAGACGGCATACGAGAAGTCGTCGTGACTGGAGTTCAGACGTGTGCTCTTCCGATCTGCTGCCGGGTATTTGTCAAA | hilD |
| AN82 | CAAGCAGAAGACGGCATACGAGAACGATTAGTGACTGGAGTTCAGACGTGTGCTCTTCCGATCTGCTGCCGGGTATTTGTCAAA | hilD |
| AN83 | CAAGCAGAAGACGGCATACGAGAACTTCGAGTGACTGGAGTTCAGACGTGTGCTCTTCCGATCTGCTGCCGGGTATTTGTCAAA | hilD |
| AN84 | CAAGCAGAAGACGGCATACGAGAATGGTCAGTGACTGGAGTTCAGACGTGTGCTCTTCCGATCTACTCCACTGTCACGCTTAAC | hilD |
| AN85 | CAAGCAGAAGACGGCATACGAGAACACTGTGTGACTGGAGTTCAGACGTGTGCTCTTCCGATCTACTCCACTGTCACGCTTAAC | leuO |
| AN86 | CAAGCAGAAGACGGCATACGAGAACTGATCGTGACTGGAGTTCAGACGTGTGCTCTTCCGATCTACTCCACTGTCACGCTTAAC | leuO |
| AN87 | CAAGCAGAAGACGGCATACGAGAATTTCACGTGACTGGAGTTCAGACGTGTGCTCTTCCGATCTACTCCACTGTCACGCTTAAC | leuO |
| AN88 | CAAGCAGAAGACGGCATACGAGAACCACTCGTGACTGGAGTTCAGACGTGTGCTCTTCCGATCTACTCCACTGTCACGCTTAAC | leuO |
| AN89 | CAAGCAGAAGACGGCATACGAGAAATCAGTGTGACTGGAGTTCAGACGTGTGCTCTTCCGATCTACTCCACTGTCACGCTTAAC | leuO |
| AN90 | CAAGCAGAAGACGGCATACGAGAAAGGAATGTGACTGGAGTTCAGACGTGTGCTCTTCCGATCTACTCCACTGTCACGCTTAAC | leuO |
| AN91 | CAAGCAGAAGACGGCATACGAGAATAGTTGGTGACTGGAGTTCAGACGTGTGCTCTTCCGATCTACTCCACTGTCACGCTTAAC | leuO |
| AN92 | CAAGCAGAAGACGGCATACGAGAAAAAATGGTGACTGGAGTTCAGACGTGTGCTCTTCCGATCTACTCCACTGTCACGCTTAAC | leuO |
| AN93 | CAAGCAGAAGACGGCATACGAGAAGTCGTCGTGACTGGAGTTCAGACGTGTGCTCTTCCGATCTACTCCACTGTCACGCTTAAC | leuO |
| AN94 | CAAGCAGAAGACGGCATACGAGAACGATTAGTGACTGGAGTTCAGACGTGTGCTCTTCCGATCTACTCCACTGTCACGCTTAAC | leuO |
| AN95 | CAAGCAGAAGACGGCATACGAGAACTTCGAGTGACTGGAGTTCAGACGTGTGCTCTTCCGATCTACTCCACTGTCACGCTTAAC | leuO |

[a] Red lettering denotes PCR priming sequences. Colored hexameric sequences denote indexes used in Illumina sequencing.
